# Confidence Read from Others’ Movements Guides Collective Decisions

**DOI:** 10.64898/2026.09.14.748905

**Authors:** Laura Schmitz, Noemi Montobbio, Mariacarla Memeo, Oussama Sabri, Bahador Bahrami, Andrea Cavallo, Stefano Panzeri, Cristina Becchio

## Abstract

A central challenge in decision-making is deciding when to trust one’s own judgement and when to rely on others. Because subjective confidence correlates with objective accuracy, a sensible strategy is to follow the choice made with higher confidence. Crucially, however, this strategy depends on accessing others’ confidence. Here, we show that humans spontaneously ‘read out’ confidence from others’ movements and use this information to guide collective decisions. Dyads performed a collective perceptual decision-making task. In each trial, one participant—the arbitrator—first made an individual decision by reaching toward one of two targets, then observed their partner doing the same, and finally made the collective decision on behalf of the dyad, confirming or revising their initial choice. When own confidence was low or high, arbitrators relied on simple heuristics—revising their choice when uncertain and confirming it when certain. At intermediate confidence levels, however, they extracted confidence from their partner’s movement kinematics to guide revision. Computational simulations demonstrated that integrating the partner’s encoded confidence improved collective accuracy. Our findings highlight the importance of bodily motion for the social transmission of confidence information in collective decision-making.

**Impact statement:** People use confidence inferred from others’ actions to revise their own choices and improve collective decision-making under uncertainty.

---

In daily life, most decisions are not made in isolation but together with, or after consulting, others (Bang & Frith, 2017; Kameda et al., 2022; Pescetelli et al., 2021; Tindale & Winget, 2019). When encountering disagreement in such *collective decisions*, individuals must assess whether and to what extent to trust others’ judgements and accordingly, whether or not to revise their own (Resulaj et al., 2009; Soll & Larrick, 2009; Stone et al., 2022). As subjective confidence correlates with objective performance (Henmon, 1911; Nelson & Narens, 1990; Peirce & Jastrow, 1884), a sensible strategy is to follow, in each case, the choice associated with the highest confidence. Following this strategy, “two heads can be better than one”: by selecting the most confident opinion in each disagreement, dyads can indeed outperform their best individual member (Bahrami et al., 2010, 2012a, 2012b; Bang et al., 2014; Fusaroli et al., 2012).

Crucially, however, this approach depends on individuals being able to access their partner’s confidence. Although substantial research has examined the ability to report and utilize one’s own confidence (Balsdon et al., 2020; Henmon, 1911; Mamassian & de Gardelle, 2022; Nelson & Narens, 1990; Peirce & Jastrow, 1884; Rahnev et al., 2020), much less is known about how others’ confidence is estimated (Bang et al., 2022) and incorporated. Previous work on collective decision-making has largely focused on settings in which dyad members explicitly communicate their confidence (Bahrami et al., 2010, 2012a, 2012b; Fusaroli et al., 2012), leaving open the question of whether this inference is restricted to the domain of language and, if not, how others’ confidence is inferred nonverbally.

One potential implicit source of confidence information is motor behavior. Accumulating evidence suggests that confidence (Dotan et al., 2018; Macerollo et al., 2015; Palmer et al., 2016; Patel et al., 2012)—as well as other mental states long considered inaccessible to others (Goldman, 2012) such as intentions (Cavallo et al., 2016; Lewkowicz et al., 2015), expectations (Grèzes et al., 2004; Runeson & Frykholm, 1983), attitudes (Manera et al., 2011), beliefs (van der Wel et al., 2014), and emotions (Dittrich et al., 1996; Pollick et al., 2001)—is reflected in subtle variations in movement patterns (Ansuini et al., 2015; Becchio et al., 2018, 2024). This raises the possibility that confidence information can be inferred from observing others’ actions (Macerollo et al., 2015; Patel et al., 2012). Yet, whether interacting decision-makers extract (‘read out’) confidence information from each other’s movements and use this information to revise their own decisions in real-time remains unknown.

To address this question, we combined kinematic analysis with computational modeling in a collective, dyadic decision-making task. On each trial, one participant—acting in the role of the arbitrator—first made an individual perceptual decision in a visual search task and expressed their choice by reaching towards one of two response options. Next, they observed their partner expressing their own choice. The arbitrator could then either confirm their initial choice or revise it when making the collective decision. Results revealed that arbitrators relied on some simple heuristics: they revised their initial choice when unsure but stuck with it when sure. When their own confidence was intermediate, however, they drew on the confidence conveyed by their partner’s actions to guide the collective decision. These results demonstrate that observers spontaneously extract confidence information from others’ movements and integrate it into the decision-making process to improve collective performance.

## Results

Fifteen dyads performed a Two-Alternative Forced Choice (2AFC) perceptual decision-making task which required the detection of a target stimulus in one of two consecutively presented viewing intervals (see schematic in Fig. 1B). To indicate their choice (interval 1 or 2), participants reached towards a response button on the left or right, respectively (see Fig. 2A). Each trial was structured such that the dyad member who acted as arbitrator first made their individual decision (Fig. 1A, left panel). Then the arbitrator observed the partner making their individual decision (Fig. 1A, middle panel). After observing the partner’s decision, the arbitrator made the collective decision on behalf of the dyad, either confirming their initial choice or revising it (Fig. 1A, right panel). Dyad members alternated, trial by trial, in the role of arbitrator. They were instructed to collaborate as a team with the goal of maximizing the dyad’s collective accuracy. No accuracy feedback was provided. After each decision, participants privately indicated how confident they felt in that decision on a scale from one to six, using a keypad.

**Fig. 1.**
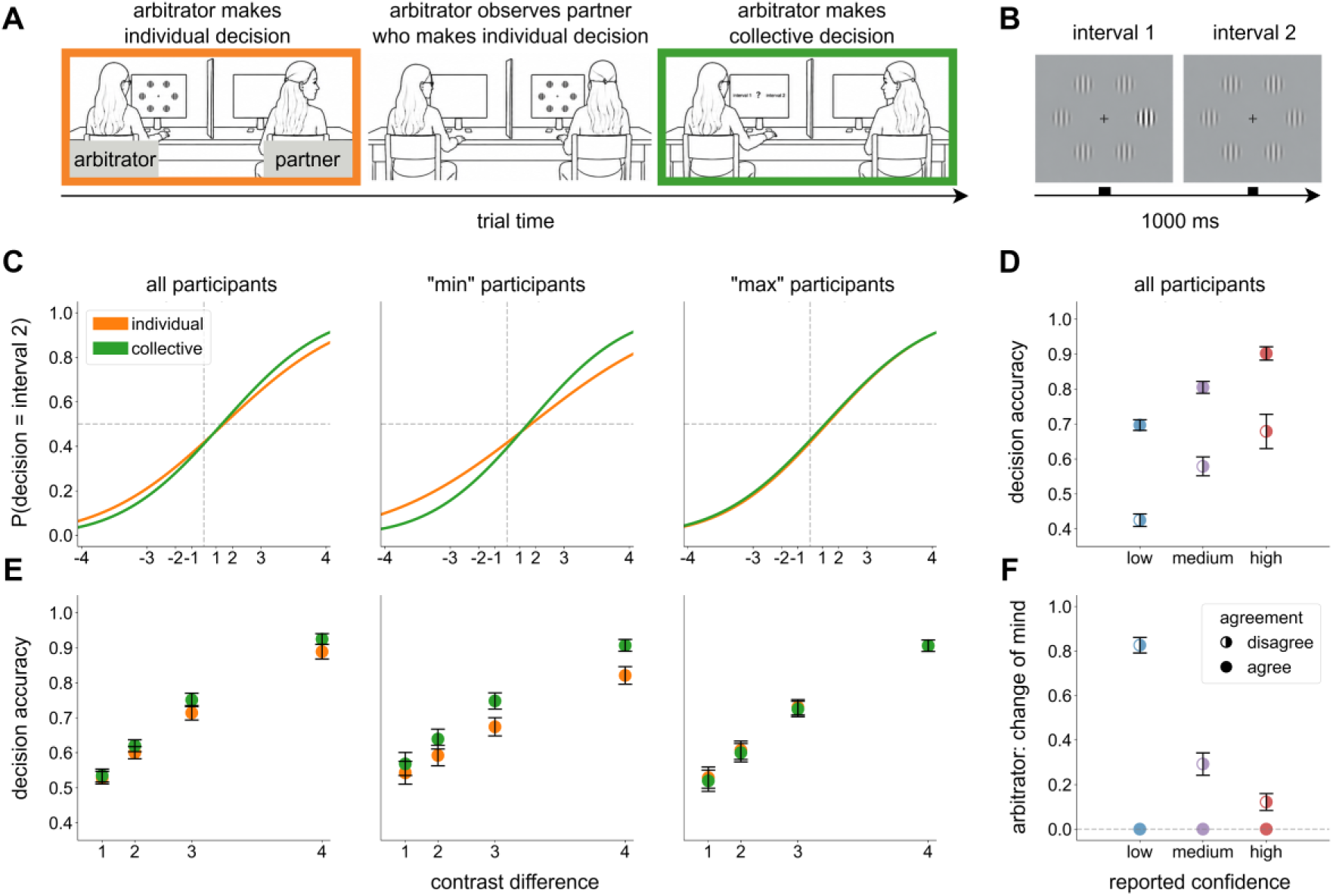
Experimental design and decision performance. (A) Setup and trial procedure. Dyad members were seated side by side in front of separate computer screens. A divider positioned between them ensured that each participant could see only the partner’s arm when it reached forward. On each trial, the designated arbitrator first made an individual perceptual decision and then observed the partner making their individual decision. After observing the partner’s decision, the arbitrator made the collective decision on behalf of the dyad. (B) Perceptual task. In a two-alternative forced-choice (2AFC) task, two stimulus intervals were presented sequentially, each containing six Gabor patches arranged in a circular array. One interval contained a target patch with higher contrast than the remaining patches. Participants indicated which interval contained the target. (C) Perceptual sensitivity. Psychometric curves represent the probability of choosing interval 2 as a function of contrast difference, with negative and positive values indicating trials in which the target occurred in interval 1 or interval 2, respectively. Slopes are shown separately for individual and collective decisions, and for all participants, as well as for the less (“min”) and more (“max”) sensitive dyad members separately. For visualization, contrast levels are labeled from 1-4 on the x-axis, corresponding to increasing contrast differences (0.015, 0.035, 0.07, 0.15). (D) Individual decision accuracy as a function of participants’ reported confidence (categorized into three levels: low, medium, high) for agreement and disagreement trials. (E) Decision accuracy as a function of contrast difference. Accuracies are shown separately for individual and collective decisions, and for all participants, as well as for the less (“min”) and more (“max”) sensitive dyad members separately. (F) Probability of change of mind as a function of the arbitrator’s reported confidence for agreement and disagreement trials. Error bars in panels D, E and F show the estimated marginal means (EMMs) +/- SEM as obtained from mixed-effects modeling.

**Fig. 2.**
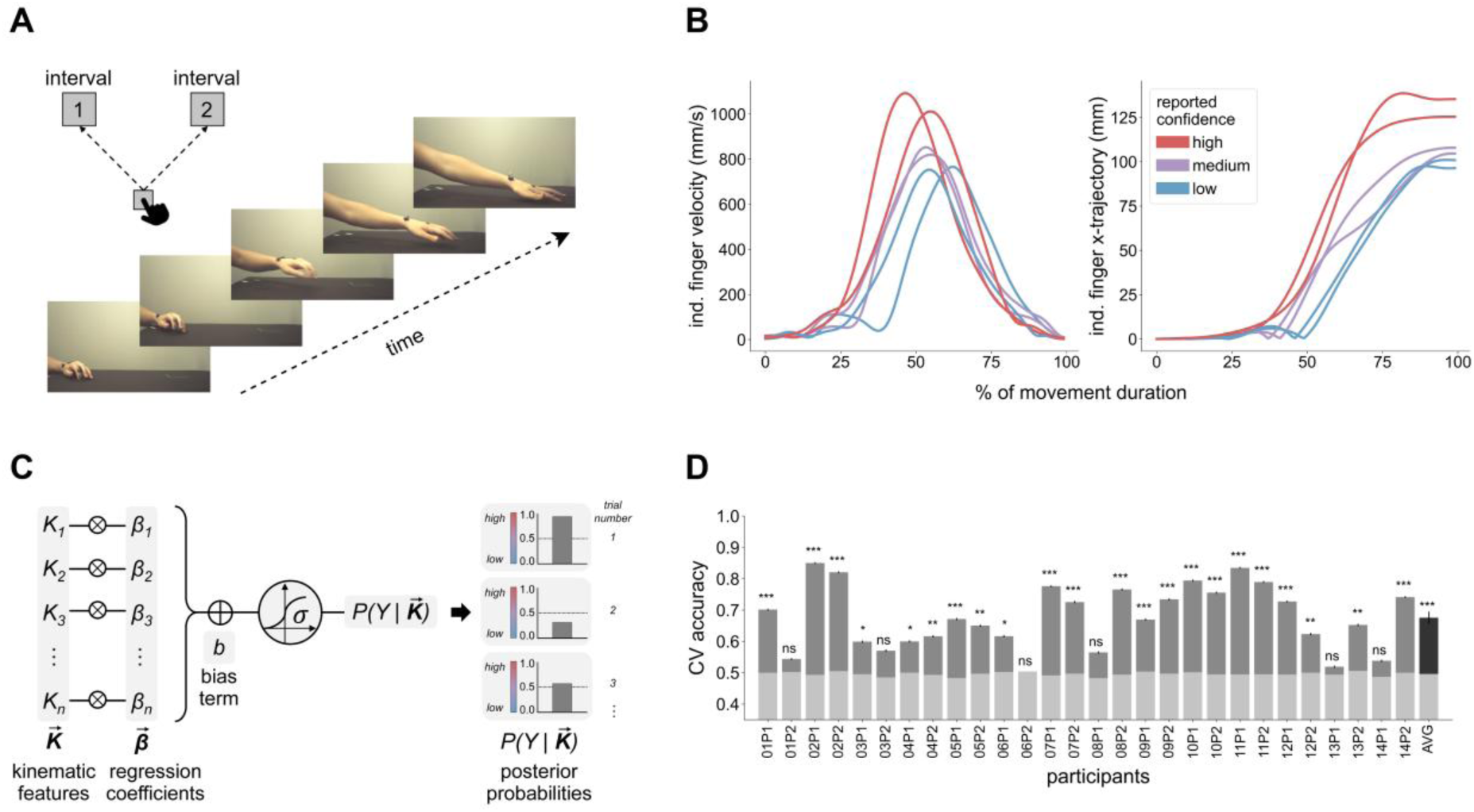
Kinematic encoding of confidence. (A) Example video frames of a reaching movement towards one of the two response buttons. For display purposes, frames are flipped over the vertical midline with respect to the original orientation. The schematic in the upper left corner represents the layout of the start and the two response buttons. (B) Time course of velocity (left panel) and x-trajectory (transverse direction, i.e., deviation from midline; right panel) of the index finger for reaching movements performed with different levels of confidence. Single lines show representative movement trajectories for one participant; colors indicate confidence level. For display and analysis purposes, x-trajectories are flipped over the midline such that all are directed towards the same side. (C) Schematic of the confidence encoding model (EM). For each choice, movement kinematics were represented as a 28-dimensional feature vector (7 kinematic variables over 4 time epochs) and entered into a logistic regression model to predict the participant’s reported confidence. Separate encoding models were trained for each participant. (D) Cross-validated model performance, quantified as the fraction of choices for which reported confidence was correctly predicted from the kinematics. Stratified K-Fold Cross-Validation (100 x 5-fold, 1000 repetitions) was applied. Model performance is shown for individual participants and averaged across participants (AVG). Bars represent mean ± SEM across folds. ns > 0.05; *p < 0.05; **p < 0.01; ***p < 0.001.

### Confidence predicts accuracy

For confidence to be useful for guiding collective decisions, it must be predictive of accuracy. As a first step, we therefore verified that participants’ reported confidence (low, medium, high) predicted the accuracy of their individual choices. Mixed-effects analysis revealed a robust relationship between confidence and accuracy, in line with previous research (Henmon, 1911; Nelson & Narens, 1990; Peirce & Jastrow, 1884). As shown in Figure 1D, higher confidence was associated with a higher likelihood of being correct, both when dyad members agreed and when they disagreed in their individual choices (i.e., relative to intermediate confidence levels, accuracy was significantly higher for high confidence levels (est = 0.431, SE = 0.207, z = 2.082, *p* = .037) and lower for low confidence levels (est = -0.621, SE = 0.122, z = -5.103, *p* < 0.001); Table S1).

### Dyads achieve a collective benefit

In perceptual decision tasks, a collective benefit occurs when the sensitivity of the collective decision exceeds that of the individual decisions (Kameda et al., 2022; Bahrami et al., 2010, 2012a; Fusaroli et al., 2012). In our design, this meant that arbitrators needed to improve upon their own initial decisions when making the collective decision after observing their partner’s action. To quantify this, following standard psychophysical methods, we constructed separate psychometric curves for individual and collective decisions. These curves describe the probability of reporting the target in interval 2 as a function of target contrast (0.015, 0.035, 0.07, 0.15—with trials in which the target occurred in interval 1 or interval 2 represented as negative and positive contrast values, respectively). The individual curve was based on trials in which participants, acting as arbitrators, made the initial individual decision (Fig. 1A, first panel), whereas the collective curve was based on the corresponding collective decision made by the same participants on those same trials after observing their partner’s action (Fig. 1A, third panel). We then used linear mixed-effects models to compare individual versus collective slopes at the population level.

Model results revealed a significant interaction between decision type and contrast level (est = 1.706, SE = 0.566, z = 3.011, *p* = 0.003, Tables S2a), reflecting steeper slopes and thus higher sensitivity for collective decisions compared to individual decisions (Fig. 1C, left panel). Analyses of decision accuracy (measured as proportion correct) confirmed this result, also showing a significant interaction between decision type and contrast level (est = 3.000, SE = 1.509, z = 1.988, *p* = 0.047, Tables S3a), reflecting higher accuracy of collective decisions relative to individual decisions at higher contrast levels (Fig. 1E, left panel).

To characterize how the collective benefit relates to individual differences in perceptual sensitivity within dyads, following Bahrami et al. (2010, 2012a, 2012b), we next classified, within each dyad, the two participants as the more and less sensitive dyad member. We then analyzed separately trials in which the less sensitive member acted as arbitrator (“min”) and trials in which the more sensitive member acted as arbitrator (“max”). This analysis revealed a significant improvement in sensitivity for less sensitive dyad members (est = 3.417, SE = 0.794, z = 4.301, *p* < 0.001; Tables S2b; Fig. 1C, middle panel) but not for more sensitive dyad members (est = −0.147, SE = 0.819, z = −0.179, *p* = 0.858; Table S2c; Fig. 1C, right panel). A similar pattern was observed for decision accuracy: less sensitive members showed accuracy gains for collective decisions (est = 4.809, SE = 2.092, z = 2.299, *p* = 0.022, Tables S3b; Fig. 1E, middle panel) whereas no significant accuracy gains were observed for more sensitive members (est = 0.243, SE = 2.252, z = 0.108, *p* = 0.914, Table S3c; Fig. 1E, right panel). Together, these results indicate that, at the population level, collective decisions selectively enhanced the performance of the less sensitive members.

### Arbitrator’s confidence and partner’s choice predict changes of mind

In our experiment, collective performance can only exceed individual performance if arbitrators revise their initial individual choices effectively. To understand how such effective “changes of mind” occurred, we examined the potential sources of information available to arbitrators.

A first potential source of information is arbitrators’ own confidence. As shown in Fig. 1D, confidence reliably predicted the accuracy of individual choices. If arbitrators relied on their own confidence to decide whether to revise their initial choices or to maintain them, the probability of a change of mind should increase as confidence decreased. This is exactly what we observed: the probability that arbitrators changed their mind was significantly higher at low confidence (0.35; est = 1.290, SE = 0.133, z = 9.684, *p* < 0.001) and lower at high confidence (0.04; est = −1.178, SE = 0.254, z = −4.633, *p* < 0.001) compared to intermediate confidence (0.13; Tables S4a). This indicates that arbitrators relied on their own confidence when deciding whether to revise their initial choice.

The second source of information is the partner’s choice. In our experiment, after making an individual decision, the arbitrator observed the partner responding and could thus determine whether the partner agreed or disagreed. Disagreement could prompt the arbitrator to reconsider their decision. If so, the probability of a change of mind should be higher when dyad members disagreed in their choices (disagreement trials) than when they agreed (agreement trials). This prediction was also confirmed: on disagreement trials, the probability that arbitrators changed their mind was 0.54 on average, with the probability of a change of mind decreasing as arbitrators’ own confidence increased. Changes of mind were significantly more likely at low confidence (0.83; est = 2.440, SE = 0.219, z = 11.163, *p* < 0.001) and less likely at high confidence (0.12; est = −1.092, SE = 0.314, z = −3.474, *p* < 0.001) relative to intermediate confidence (0.29; see Tables S4b and Fig. 1F). In contrast, on agreement trials, no changes of mind occurred at any level of confidence, indicating a complete absence of revision when the partner’s choice aligned with the arbitrator’s initial choice (Fig. 1F). This suggests that arbitrators took into account their partner’s choice and this influenced integration of own confidence on decision revision.

### Confidence is encoded in individuals’ movement kinematics

So far, we have considered two explicit sources of information immediately available to arbitrators: their own confidence and the partner’s choice. A third, implicit source is the partner’s confidence potentially conveyed through movement kinematics. For this information to influence changes of mind, two conditions must be satisfied: first, confidence must be encoded in movement kinematics at the single-choice level; second, arbitrators must extract this information from the partner’s observed movement and integrate it when making the collective decision.

As a first step, we therefore tested whether movement kinematics predict the partner’s confidence (privately reported, using a keypad, on a scale from 1-6 after each choice) on a choice-by-choice basis. To do so, we adapted the recently developed kinematic coding framework (Becchio et al., 2024). We represented the kinematics of each choice as a 28-dimensional feature vector (7 kinematic variables over 4 time epochs) and then fitted a logistic regression model to predict the confidence reported for a given choice from the movement kinematics of that choice (Fig. 2C). The resulting posterior probability provides a measure of single-choice confidence encoding (Becchio et al., 2024). We trained an individual ‘encoding model’ for each participant and evaluated model performance as the fraction of choices for which reported confidence was correctly predicted by the model. Across choices and participants, average model performance reached 67% (95% CI [64, 71]) (Fig. 2D; rightmost dark bar; see Table S5 for details on model performance). These results indicate that movement patterns reliably encoded confidence information at the single-choice level (see Fig. 2B for an illustration).

### Partner’s confidence predicts changes of mind

Having established that movement kinematics encode confidence information, we next assessed whether arbitrators extracted this information from their partner’s actions and integrated it when revising their initial choices on disagreement trials. Previous studies using forced-choice paradigms have shown that naïve participants can read information encoded in movement patterns when explicitly instructed to do so (Cavallo et al., 2016; Montobbio et al., 2022; Patel et al., 2012; Patri et al., 2020; Scaliti et al., 2023). In our design, however, arbitrators were not instructed to infer their partner’s confidence, raising the question of whether such information would nevertheless be incorporated during decision revision.

Intuitively, relying on the partner’s confidence should be most beneficial when arbitrators’ own confidence provides limited guidance, that is, when arbitrators’ own confidence was in an intermediate range. When own confidence is either high or low, arbitrators can rely on a simple heuristic: consistently sticking to their own response when confidence is high (no change of mind) and adopting the partner’s response when own confidence is low (change of mind). At intermediate confidence, however, this heuristic provides no clear guidance—change or no change? In such cases, arbitrators may engage in more cognitively demanding strategies and incorporate their partner’s encoded confidence into the decision process.

We tested this idea by comparing a series of mixed-effects models that differed in whether and how they accounted for the partner’s confidence. All models included arbitrator’s own confidence as a predictor. We began with a model that did not contain any information about the partner’s confidence (Table S6a). We then added partner’s encoded confidence as an additional predictor (Table S6c). Finally, we tested whether arbitrators selectively rely on their partner’s confidence depending on their own confidence by fitting a piece-wise model that allowed the effect of partner’s encoded confidence to vary across the three levels of arbitrator’s own confidence (Table S6d).

Model comparison showed that this piece-wise model provided the best fit to participants’ choices, even after accounting for model complexity (Table S6e). Fixed-effects results confirmed a strong, negative effect of arbitrators’ own confidence (est = −1.762, SE = 0.152, z = −11.600, *p* < 0.001, Table S6d), indicating that the probability of a change of mind decreased as the arbitrator’s own confidence increased (Fig. 3A; also compare Fig. 1F). In addition, the model revealed a selective effect of partner’s encoded confidence that depended on the arbitrator’s own confidence: when arbitrator’s own confidence was intermediate, the partner’s confidence predicted the probability of a change of mind (est = 0.593, SE = 0.172, z = 3.450, *p* < 0.001), with arbitrators being more likely to change when the partner’s action conveyed higher confidence (see Fig. 3A, purple slope). By contrast, partner’s encoded confidence had no significant influence when arbitrator’s own confidence was low (est = 0.045, SE = 0.141, z = 0.322, *p* = 0.748) or high (est = −0.071, SE = 0.392, z = −0.182, *p* = 0.856); see Table S6d.

**Fig. 3.**
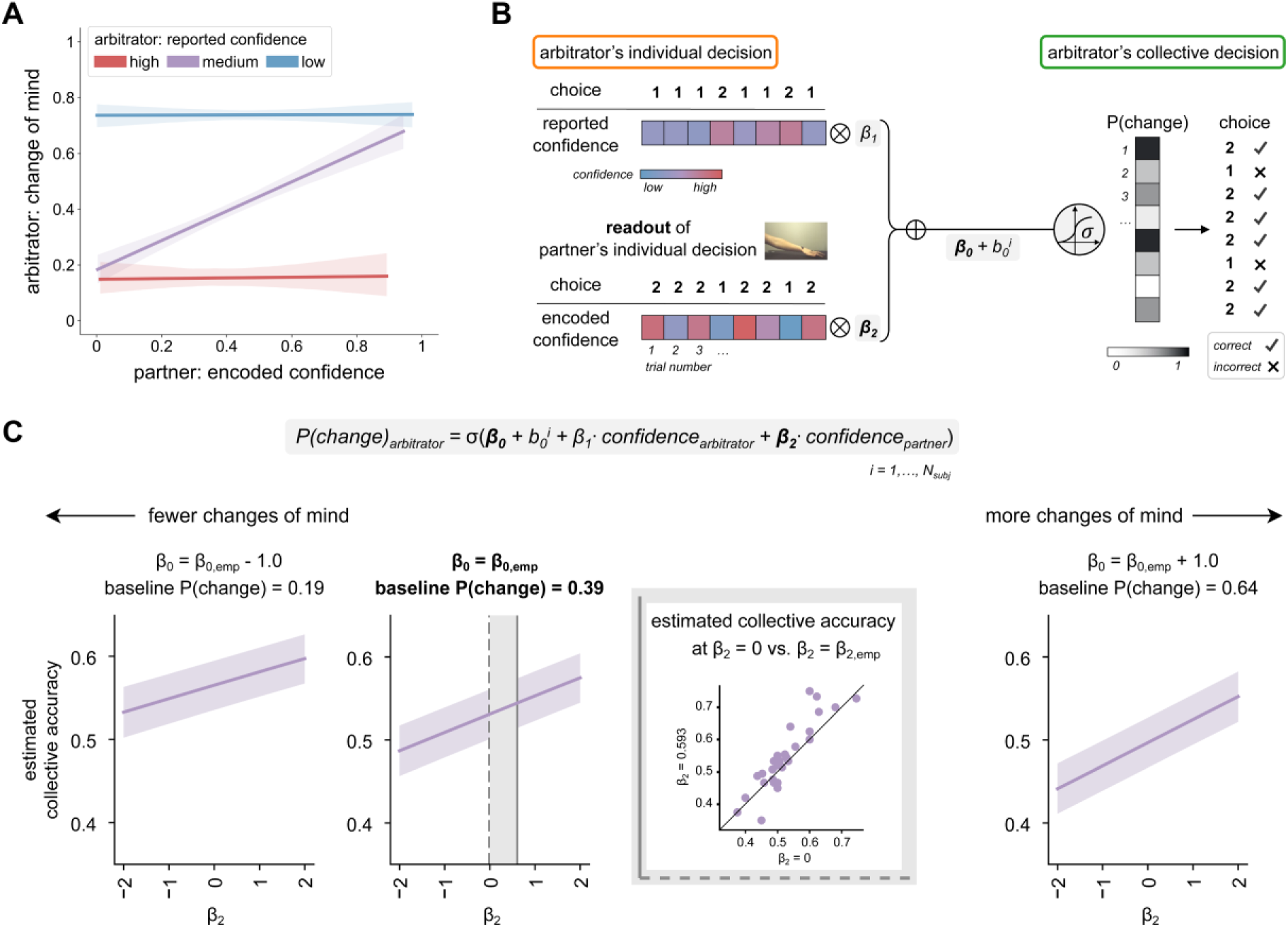
Partner’s confidence shapes changes of mind and collective accuracy. (A) Probability of a change of mind as a function of the arbitrator’s reported confidence (classified into high, medium, low) and the partner’s encoded confidence. Analysis includes disagreement trials only. (B) Schematic of the computational model. The change-of-mind probability of the arbitrator is defined by a logistic function that includes the baseline change-of-mind probability (β_0_), the effect of the arbitrator’s own confidence (β_1_), the effect of the partner’s encoded confidence (β_2_), and a subject-level random intercept (*b*^*i*^_0_). In simulations, β_0_ and β_2_ were systematically varied, while β_1_ and b^i^_0_ were held fixed at their empirically estimated values. (C) Estimated collective accuracy as a function of the strength of integration of the partner’s encoded confidence (β_2_), shown separately for different baseline tendencies to change (β_0_). The central panel corresponds to the empirically observed baseline tendency to change (β_0_ = β_0,emp_; P(change) = .39). The continuous vertical line marks the empirically observed level of integration of partner’s confidence (β_2_ = 0.593), whereas the dashed line marks a counterfactual zero-integration scenario (β_2_ = 0), in which the partner’s confidence does not contribute to the probability of change. The inset scatter plot shows a paired comparison of the estimated collective accuracy under the empirical integration level (β_2_ = β_2,emp_ = 0.593) versus the counterfactual zero-integration scenario (β_2_ = 0); points above the identity line indicate higher collective accuracy when the partner’s confidence information is incorporated.

Together, these findings indicate that partner’s *encoded* confidence was selectively integrated at intermediate levels of arbitrator’s own confidence during decision revision. As a control analysis, we tested whether the partner’s *reported* confidence predicted changes of mind. In contrast to encoded confidence, reported confidence did not significantly predict changes of mind (est = 0.130, SE = 0.107, z = 1.220, *p* = 0.223; Table S6b), demonstrating that partner’s confidence influenced decision revision only insofar as it was encoded in movement kinematics and thus accessible to the arbitrator.

### Integrating partner’s confidence improves collective decisions

The results above indicate that, at intermediate levels of own confidence, arbitrators integrated their partner’s encoded confidence when revising their initial choices. Together with the finding that confidence predicts accuracy (Fig. 1D), this implies that incorporating the partner’s confidence improves collective decision accuracy. To illustrate this in our data and to obtain an estimate of the effect, we developed a simple computational model. This model enabled us to systematically vary the strength of this integration at intermediate levels of arbitrator’s own confidence and assess its effects on collective performance, quantified as collective accuracy.

Specifically, the model (schematized in Fig 3B) computed collective accuracy based on the simulated probability that the arbitrator changes their mind. This probability was defined by a logistic function with four components, matching our piece-wise model introduced above (Table S6e): the baseline change-of-mind probability (*β*_0_), the sensitivity of the change probability to the arbitrator’s own reported confidence (*β*_1_), the sensitivity of the change probability to the partner’s encoded confidence (*β*_2_ ; see purple curve in Fig. 3A), and the subject-level deviation from the intercept (*b*^*i*^_0_) Simulated probabilities of change of mind were then used to generate collective decisions by combining the arbitrator’s initial response with the simulated revision (change or no change). Collective accuracy was computed as the proportion of trials in which the simulated collective decision was correct.

To anchor the simulations to the experimentally observed scenario, we first evaluated model performance using the empirical parameters obtained from our mixed-effects analysis reported above (*β*_0_ = *β*_0,*emp*_, corresponding to a baseline probability of change of mind of 0.39, and *β*_2_ = *β*_2,*emp*_ = 0.593). Under these parameters, estimated collective accuracy closely matched the empirical collective accuracy (central panel of Fig. 3C), indicating that the model captured the main determinants of collective performance and provided a reliable estimate of the empirical data.

To quantify the effect of integrating encoded confidence under the empirical baseline tendency of change, we then systematically varied the strength of integration (*β*_2_) across a wide range of values (from –2.0 to 2.0), while holding *β*_0_ fixed at its empirical estimate. Increasing *β*_2_ led to a significant increase in simulated collective accuracy (slope = 0.082, SE = 0.004, *p* <0. 0001; see purple curve in central panel of Fig. 3C and Table S7a), indicating that greater sensitivity to the partner’s encoded confidence systematically improves collective performance. The inset scatter plot shows the paired comparison of the estimated collective accuracy under the empirical level of integration (*β*_2_ = *β*_2,*emp*_ = 0.593; continuous vertical line in the central panel of Fig. 3C) with a zero-integration counterfactual scenario (*β*_2_ = 0; dashed vertical line), in which the partner’s confidence did not contribute to the probability of change. Most points lie above the identity line, indicating that incorporating the partner’s confidence information systematically improves collective performance despite variability across simulations (paired comparison, *p* < 0.001).

To assess whether this benefit generalized beyond the empirical baseline tendency of change, we repeated the simulations while varying *β*_0_ (from *β*_0,*emp*_ – 2.0 to *β*_0,*emp*_ + 2.0; see left and right panels of Fig. 3C for *β*_0,*emp*_ – 1.0 and *β*_0,*emp*_ + 1.0, respectively). Across all baseline levels, simulated collective accuracy increased as a function of *β*_2_ (see Table S7b). This indicates that integrating the partner’s encoded confidence robustly benefited collective performance across a wide parameter space and regardless of the assumed baseline tendencies of changing one’s mind. All trends described above remained consistent when generating single-trial collective decisions stochastically (see Tables S7c-d).

## Discussion

A central challenge in collective decision-making is determining when to trust one’s own judgment and when to rely on others. This challenge is especially pronounced when one’s own confidence provides limited guidance. Here we show that in such cases, confidence inferred from others’ actions can guide decision revision and yield a measurable collective benefit. We tested this in a perceptual decision-making task where dyad members took turns in the role of arbitrator: they first made an individual choice by reaching toward a target, then observed their partner doing the same, and finally made the collective decision on behalf of the dyad, confirming or revising their initial choice.

When dyad members agreed in their initial individual choices, no changes of mind occurred, regardless of the arbitrators’ confidence levels. This absence of revision under consensus demonstrates the strong social influence (Moussaïd et al., 2013) of partner agreement, effectively overriding even low confidence in one’s own initial decision.

When dyad members disagreed, however, revisions depended on arbitrators’ own confidence and on their partner’s confidence. Arbitrators maintained their initial choice when own confidence was high and followed their partner’s choice when own confidence was low. This is in line with related prior work on advice seeking, which shows that confidence predicts advice requests: individuals seek less advice when they are more confident in their initial decisions (Pescetelli et al., 2021). At intermediate levels of own confidence, however, arbitrators drew on their partner’s confidence, as expressed through their movements, to guide revision. Combining kinematic analysis with mixed-effects model analysis enabled us to show that revisions were predicted by confidence encoded in the partner’s movement at the single-choice level. Computational modeling further showed that integrating partner’s encoded confidence increased collective accuracy across a broad range of baseline revision tendencies, suggesting that kinematic readout of confidence can serve as a mechanism to improve collective decisions.

Previous studies investing kinematic readout have typically relied on passive observation paradigms, where participants viewed others’ actions offline and made forced-choice judgments about the encoded information (Cavallo et al., 2016; Lewkowicz et al., 2015; Montobbio et al., 2022; Patel et al., 2012; Patri et al., 2020; Scaliti et al., 2023). Our results advance this work along three dimensions. First, they demonstrate that confidence information can be inferred from others’ movements not only during offline observation (Macerollo et al., 2015; Palmer et al., 2016; Patel et al., 2012) but also during real-time social interaction (Pezzulo et al., 2013, 2019; Sebanz et al., 2006; Sebanz & Knoblich, 2021; Vesper et al., 2017). This extension is theoretically significant as mechanisms supporting social interaction have been proposed to fundamentally differ from those supporting social observation, where individuals observe social stimuli without the opportunity for interaction (Redcay & Schilbach, 2019; Schilbach et al., 2013). During social interaction, individuals do not merely observe others’ behavior, but prepare, update, and coordinate their own responses in relation to it (Konvalinka et al., 2010; Sebanz et al., 2006; Sebanz & Knoblich, 2021). In this context, kinematic readout ceases to be a mere source of information for third-person inference and instead becomes part of a first-person control process that shapes ongoing decision-making.

Second, our results show that confidence information is used online to guide decision revision even when participants are given the opportunity to ignore each other’s actions. Previous work has shown that in settings requiring consensus, decision outcomes tend to be dominated by group members whose actions convey higher confidence (Coucke et al., 2024). Our findings extend this view by demonstrating that even in settings where consensus is not enforced, collective decisions are influenced by confidence implicitly encoded in others’ movement kinematics. Notably, this effect was not explained by the partner’s explicitly reported confidence, indicating that the critical signal guiding revision was encoded in the kinematics of the action. This highlights the functional significance of movement kinematics for calibrating the weight assigned to social information, allowing individuals to adjust the influence of others’ input in proportion to its inferred reliability.

Third, our results reveal that the integration of others’ confidence is calibrated to the arbitrator’s own confidence. The fact that participants incorporated their partner’s encoded confidence selectively at intermediate levels of their own confidence highlights a flexible, uncertainty-driven, metacognitive mechanism for integrating social information in collective decision-making. This selective use of information is consistent with the broader view that humans balance accuracy and complexity, recruiting more sophisticated decision-making strategies as a function of uncertainty (Pescetelli et al., 2021; Tavoni et al., 2022). When uncertainty is low or high, complex strategies offer little advantage over simpler heuristics. At intermediate levels of uncertainty, however, more demanding strategies become worthwhile despite their potential cognitive cost. In our task, this trade-off is evident: participants relied on simple heuristics when their own confidence was low or high but engaged in more demanding strategies when their own confidence was at intermediate levels. This pattern of results is consistent with the notion that metacognitive monitoring governs not only evaluations of one’s own cognitive state but also the integration of information about others’ cognitive states (Bang et al., 2022).

Together, these findings provide direct evidence for adaptive transmission and usage of confidence in social decision contexts. They show that people can flexibly recruit complex inferential mechanisms based on movement information when self-confidence alone is insufficient, and that doing so yields a quantifiable collective advantage. This proposes a principled mechanism through which sensorimotor communication supports collective intelligence (Burton et al., 2024; Galton, 1907; Surowiecki et al., 2004): confidence dynamically coordinates information flow across individuals, enhancing group-level performance without explicit signaling or verbal exchange.

## Materials and Methods

### Participants

Thirty-two participants were recruited from the participant pool of the Istituto Italiano di Tecnologia in Genova, Italy. All participants fulfilled the following inclusion criteria: aged between 18 and 40 years, normal or corrected-to-normal vision, right-handed, Italian-speaker, no neurological or psychiatric disease history. For each experimental session, two individuals were invited to participate together as a dyad. Dyad members did not know each other. Dyads were composed of same-sex individuals, i.e., female-female and male-male dyads.

One dyad was excluded from all analyses because the participants did not follow the experimenter’s instructions. Due to technical error, kinematic data from another dyad were not available. Thus, the final sample included N=15 dyads for behavioral analyses (10 female, 5 male; age range = 19-37 years, mean = 25.03, SD = 4.33) and N=14 dyads for kinematic analyses (9 female, 5 male).

The study protocol and analysis plan were preregistered (https://doi.org/10.17605/OSF.IO/TXCW6) via OSF. A target number of 30 individuals (15 dyads) was specified a priori following studies by Bahrami and colleagues using a comparable perceptual decision-making paradigms (Bahrami et al., 2010, 2012a, 2012b, Bang et al., 2014). For kinematic encoding of confidence, separate models were fit for individual participants. The critical factor for these analyses was thus the number of trials contributed by each participant rather than the overall number of participants. Pilot testing confirmed that the available trials per participant (n = 160) were sufficient to support reliable regression analyses at the single-subject level.

Written informed consent was obtained from all participants prior to participation and participants received monetary compensation. The study was conducted in accordance with the Declaration of Helsinki (World Medical Association, 2013) and approved by the local Ethics Committee CER Liguria (Registry No. 192/2015, DB ID 2543).

### Experimental design and procedures

#### Apparatus

Two participants were seated next to each other at the long side of a table (1 × 2 m), with a distance of about one meter between them (Fig. 1A and Fig. S1A). Half-way, a partition (1.50 × 2 m) was positioned between them to prevent visual access, ensuring that each participant could see only the partner’s arm when it reached forward. The table surface in front of each participant was covered with a two-layer plastic panel (62 × 64 cm) with a black cardboard surface (Fig. S1B). Embedded in the panel were three square-shaped sensors which were sensitive to both press and release actions. There were two large sensors (8 × 8 cm) and one small sensor (4 × 4 cm). The small sensor was positioned centrally in front of the participant, on the panel’s midline and 12 cm away from its front edge. This sensor served as the start position for participants’ actions. The two large sensors were placed 36 cm away from the start position, offset by 12 cm to the right and left of the panel’s midline, respectively. These sensors served as response positions. Above the response positions, at the far edge of the panel, two small white labels were placed; the left label was numbered “1°” and the right was numbered “2°”. These numbers specified how the response positions were assigned to the binary response options: If participants chose interval 1, they should reach to the left position; if they chose interval 2, they should reach to the right position (see *Trial structure* for task details). The spatial layout of the three positions/sensors was such that participants could comfortably reach with their right hand from the start position towards either of the response positions. Next to the start position, two short parallel lines on the panel informed participants about where to place their right forearm at the beginning of each trial while placing their index finger on the start sensor. At the far edge of the table, behind the panel, two 22-inch computer screens (resolution: 1,280 × 1,024; refresh rate: 60 Hz) were positioned, one in front of each participant. Each screen was divided in half vertically such that one half was visible while the other half was covered by a black cardboard. The right half of the left screen was covered and the left half of the right screen was covered (Fig. S1A). This was done so that different information could be displayed to the two participants while both screens were connected to the same computer. The screens were positioned such that the visible half was centrally aligned with the respective start position. Participants could see only their own screen; visual access to the other participant’s screen was blocked by a black partition that was placed next to the screens (Fig. S1B). Finally, each participant was equipped with a small wireless numeric keypad. The keypad was covered with black cardboard such that only three vertically aligned keys were accessible. These three keys were marked with white tape: the middle key was fully covered with a white square while the tape on the outer two keys was shaped like an arrow, with one pointing up and the other pointing down (Fig. S1B). The keypad served for participants to indicate how confident they felt in their decision. This was done by moving a slider on a scale displayed on the screen (Fig. S1C; see *Trial structure* for details). During the experiment, the keypad rested on participants’ left knee under the table so that participants could comfortably use it with their left hand while the right hand remained on the table.

#### Task and stimuli

Participants performed a contrast discrimination task using a Two-Alternative Forced Choice (2AFC) design. On each trial, stimuli were presented sequentially in two consecutive viewing intervals. Each interval displayed six vertically oriented Gabor patches arranged in a circle (see schematic in Fig. 1B). All patches had a baseline contrast of 0.10. On every trial, one randomly selected patch in one of the two intervals was increased by one of four possible contrast increments (0.015, 0.035, 0.07, 0.15). This higher-contrast patch was the target. Participants were instructed to indicate the interval that contained the target by making a reaching movement to the left (interval 1) or right (interval 2) response position (see *Apparatus*). Target location was chosen randomly in each trial. The order of target contrast and interval was randomized across each of the two blocks of the experiment while ensuring that an equal number of the four target contrasts was presented per interval and per block. Dyad members always viewed identical stimuli within each trial.

#### Experimental structure

For each session, two participants were invited to perform the contrast discrimination task together as a dyad. On every trial, one participant acted as the arbitrator, and the other as the partner. Each trial had three phases (see Fig. 1A): First, the arbitrator made their individual perceptual decision. Second, the partner made their individual decision while the arbitrator observed. Finally, the arbitrator made the collective decision on behalf of the dyad, either confirming or revising their initial choice. Choices were indicated through reaching movements. Specifically, participants were asked to reach to the left response position if they thought the target occurred in interval 1, and to reach to the right response position if they thought the target occurred in interval 2 (as described above, see Apparatus). The roles of arbitrator and partner alternated trial by trial.

The experiment was structured in two blocks with 80 trials each, with short breaks after every 40 trials. During the breaks, participants received feedback about the collective accuracy (i.e., the average accuracy of the collective decisions so far). No individual feedback was provided. Before the start of the experiment, participants received detailed instructions. The experimenter emphasized that the core idea of the experiment was the “team aspect” and that the collective decision was more important than the individual decisions, since only the team score (i.e., the accuracy of the collective decisions) mattered in the end. To increase participants’ motivation, they were informed that the best performing dyad would receive a small prize at the end. Participants completed 16 practice trials to familiarize themselves with the task and procedure. The first 10 practice trials contained targets with the highest contrast (0.15) to facilitate detection, followed by six trials sampling the remaining three contrast levels (0.035, 0.07, 0.15). The experiment lasted 2h 15min in total and was implemented in MATLAB (The MathWorks Inc., 2022) using the Cogent 2000 toolbox (https://github.com/lnnrtwttkhn/Cogent2000).

#### Trial structure

At the beginning of each trial, an on-screen message informed participants who would act as arbitrator. After confirming readiness, the experimenter manually initiated the trial.

The arbitrator placed their index finger on the start sensor, while the partner was instructed to turn away (to prevent her from observing the arbitrator’s decision). After a 2,000 ms delay, a black fixation cross appeared in the center of the screen for a variable period between 500 and 1,500 ms, randomly drawn from a uniform distribution. The task stimuli were then presented sequentially in two intervals, each displayed for 85 ms and separated by an inter-stimulus interval of 1,000 ms. Following a 500 ms delay, a decision prompt appeared, cueing the arbitrator to indicate their judgment by reaching left (if the target was perceived in interval 1) or right (if the target was perceived in interval 2). Following this individual decision, the arbitrator rated their confidence on a six-level discrete scale (1 = very little confidence “pochissimo”; 6 = fully confident “moltissimo”) using a slider controlled with arrow keys on a handheld keypad (see *Apparatus*). Confidence ratings were always private and not accessible to the partner. A brief auditory signal then cued the partner to turn back and begin their turn. The partner completed the same sequence of fixation, stimulus presentation, individual reach decision, and private confidence rating. The arbitrator observed the partner’s reaching movement. After both individual phases, the two individual decisions were displayed simultaneously on the screen, informing participants whether they agreed or disagreed. The arbitrator was then instructed to return their finger to the start position. After a variable delay (750-1,250 ms), without repeating stimulus presentation, the decision prompt reappeared and the arbitrator made the final collective decision via a reaching movement while the partner observed. The arbitrator subsequently rated their confidence in this collective choice, concluding the trial.

#### Kinematic data acquisition and preprocessing

Participants’ reaching movements were tracked using a near-infrared camera motion capture system with nine optical cameras (frame rate: 100 Hz; Vicon Nexus v.2.10.3) and simultaneously filmed from a lateral viewpoint using a video camera fully synchronized with the optical cameras (frame rate: 50 Hz; resolution: 1,920×1,080; Vicon Vue). Participants’ right hand was outfitted with seven retroreflective hemispheric markers (6.4 mm in diameter). Markers were placed on the tip of the index finger, the distal interphalangeal joint of the index finger, the metacarpophalangeal joint (knuckle) of the index finger and of the little finger, and on the styloid processes of the radius and the ulna (wrist). Below, markers on the tip of the index and on the ulnar region of the wrist are referred to as “index marker” and “wrist marker”, respectively. An additional marker was placed on either the back of the forearm (participant seated on the left) or on the hand dorsum (participant seated on the right), for participant identification in the motion capture software. Each trial was individually inspected for correct marker identification and then run through a low-pass, 4^th^ order Butterworth filter with a 6-Hz cutoff. In total, 8% of trials were discarded from the analysis because of technical error. A MATLAB custom script was used to compute the following seven kinematic variables of interest:

● wrist velocity (WV), defined as the module of the three-dimensional velocity of the wrist marker (mm/s);
● wrist acceleration (WA), defined as the rate of change of wrist velocity (mm/s^2^);
● wrist height (WZ), defined as the z-component of the wrist marker (mm);
● index velocity (IV), defined as the module of the three-dimensional velocity of the index marker (mm/s);
● index acceleration (IA), defined as the rate of change of index velocity (mm/s^2^);
● index height (IZ), defined as the z-component of the index marker (mm);
● index horizontal trajectory (IX), defined as the x-component (transverse direction) of the index marker (mm).

Each variable was computed for the reaching phase of the movement, from movement onset to movement offset. Onset was defined either as the moment of start sensor release or, if this occurred earlier, the time point when wrist or index velocity first crossed a 20 mm/s-threshold (whichever marker passed this threshold first). Offset was defined as the moment of response sensor press. In order to make movements comparable between trials and participants, all kinematic variables were expressed with respect to normalized (%) rather than absolute movement duration.

### Quantification and statistical analysis

Analyses were conducted with custom scripts written in Python (Anaconda Inc., 2016; version 3.12.4) and R (R Core Team, 2021; version 2022.07).

#### Confidence-accuracy relationship

To assess whether participants’ confidence ratings predicted the accuracy of their individual decisions in the 2AFC task, we fit a generalized linear mixed effects model (GLMM) with a binomial family and a ‘logit’ link function. We considered single-choice accuracy (correct vs. incorrect) as dependent variable, confidence (discrete values from 1-6) as fixed effect, and participant as random intercept.

#### Psychometric curves

We used participants’ choices in the 2AFC task (interval 1 or interval 2) to construct psychometric curves representing the probability of choosing interval 2 as a function of contrast level—with trials in which the target occurred in interval 1 or interval 2 represented as negative and positive contrast values, respectively (see Fig. 1C). To evaluate whether psychometric slopes differed between individual and collective decisions, we fit a GLMM with a binomial family and a ‘probit’ link function^9^. We considered choice (interval 1 or interval 2) as dependent variable, decision type and contrast level and their interaction as fixed effects, and participant as random effect, with both random intercepts and random slopes for contrast level (on modeling psychophysical data with GLMMs, see Moscatelli et al., 2012).

We computed three different versions of this model: one including all participants’ data (n=30), one including only data from the more sensitive dyad members (“max”, n=15), and one including only data from the less sensitive dyad members (“min”, n=15). To determine, for each dyad, which member was more/less sensitive in the perceptual task, we first modeled participants’ individual decisions as a function of contrast level: we used choice as dependent variable, contrast level as fixed effect, and participant as random effect (random intercept and random slope of contrast level). The estimated individual slopes were then used to determine the max/min member per dyad, based on whose slope was steeper.

#### Accuracy

To assess the significance of decision type (individual vs. collective decision) and contrast level (0.015, 0.035, 0.07, 0.15) on the probability of making the correct choice, we fit a GLMM with a binomial family and a ‘logit’ link function. We considered single-choice accuracy (correct vs. incorrect) as dependent variable, decision type and contrast level and their interaction as fixed effects, and participant as random effect (random intercept and random slope of contrast level).

### Kinematic encoding of confidence information

#### Single-choice kinematic vector

To characterize the kinematics of each reaching movement, we averaged the seven kinematic variables of interest over four time epochs of 25% of the normalized movement duration (0-25%, 25-50%, 50-75%, and 75-100%). Each choice was thus represented as a 28-dimensional kinematic vector (henceforth ‘single-choice kinematic vector’) defined by 7 kinematic variables over the 4 time epochs (28 kinematic features). We verified that increasing the number of time epochs (to 5, 6, 8 or 10) did not yield significant improvements in encoding model performance (*p* > 0.1 for all comparisons).

#### Kinematic encoding model

To quantify the kinematic encoding of confidence information, we trained a logistic regression model (kinematic encoding model) to estimate the probability that a given choice (i.e., a single reaching movement) was performed with high confidence as a sigmoidal function of the single-choice kinematic vector (see Fig. 2C).

High-confidence and low-confidence choices were defined based on a median split of each participant’s self-reported confidence ratings. The logistic regression model computed the posterior probability of ‘high confidence’ for each choice as a sigmoidal transformation of a weighted sum of the components of the single-choice kinematic vector *K→*, as expressed by the following equation:

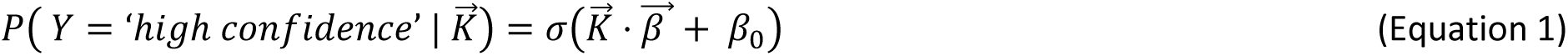

where *σ* is the sigmoid function, β→ is the vector containing the regression coefficients (weights) for each kinematic feature, *β*_0_ is a kinematic-independent bias term, and *Y* is the binary response variable predicting the confidence class (0 for low confidence; 1 for high confidence). Specifically, for each single-choice kinematic vector *K*^→^, the confidence class *Y* was predicted by computing the argmax over *Y* of *P*(*K→*); see Equation 1 and Fig. 2C.

Logistic regression was implemented using the *scikit-learn* Python library (Pedregosa et al., 2011). To correct for the slight imbalance between the two confidence classes (lower confidence ratings were more frequent than higher ratings, see Fig. S2), we specified the parameter *class_weight* as balanced, to automatically adjust weights inversely proportional to class frequencies (Thölke et al., 2023).

#### Training of logistic regression models

Encoding models were trained separately for each participant. Prior to model training, we standardized (z-scored) each participant’s kinematic vectors to ensure that predictors with larger ranges of values did not disproportionately influence the results. Encoding models were trained using L^2^ regularization. The hyperparameter *C* (commonly also referred to as *λ*), which controls the strength of the regularization term, was tuned using a nested stratified 5-fold cross-validation procedure, designed to identify a value that generalized well across participants. The two layers of cross-validation involved an outer loop for assessing the model’s performance and an inner loop for tuning the hyperparameter *C* (using a grid search over *C* values logarithmically spaced from 10^−4^ to 10^4^). This nested cross-validation procedure was run separately for each participant. We retained the value *C*=0.1, which maximized the mean cross-validated performance across participants.

Encoding models were trained on each participant’s individual choices (160 trials). These included 80 trials in which a participant took the first individual decision in the role of ‘arbitrator’ and 80 trials in which a participant took the second individual decision in the role of ‘partner’ (see *Experimental structure*). To assess robustness, we verified that model performance did not differ significantly when models were trained on only one of the two decision types (either first or second individual decisions).

#### Quantification of model performance

We evaluated the performance of the encoding models using repeated stratified 5-fold cross-validation with 100 random splits (Kim, 2009). Stratification ensured that each fold and random split preserved the original proportion of low- and high-confidence choices observed in the full dataset.

Model performance was quantified as balanced accuracy (Thölke et al., 2023), defined as the average of true positive (sensitivity) and true negative (specificity) classification rates. Sensitivity reflects the fraction of high-confidence choices correctly classified as high-confidence; specificity reflects the fraction of low-confidence choices correctly classified as low-confidence. Balanced accuracy is particularly suitable for imbalanced datasets, such as ours, where low-confidence choices (52%) slightly outnumbered high-confidence choices (48%). Classification performance reached 85% for some participants, with an average performance of 67% across all participants (Fig. 2D).

To assess statistical significance of encoding model performance, we used a permutation test approach. First, participant-level significance was determined by shuffling confidence labels (‘high’ vs. ‘low’) across trials (1000 times), retraining and evaluating the model for each permutation, and comparing the original performance to this null distribution. Second, significance across participants was assessed by calculating, for each permutation (from the first step), the mean balanced accuracy across participants. These values formed a null distribution of average balanced accuracy, which we then compared to the average balanced accuracy from the original (unshuffled) models.

For both steps, *p*-values were computed as the proportion of permuted balanced accuracies that were greater than or equal to the original (unshuffled) model’s balanced accuracy. Statistical significance was determined by comparing the computed *p*-value to the threshold 0.05.

### Dependence of changes of mind on arbitrator’s own confidence and partner’s confidence

We used GLMMs with a binomial family and a ‘logit’ link function to assess the dependence of changes of mind on the arbitrator’s own confidence and the partner’s confidence. Change of mind was defined as a collective decision in which the arbitrator changed their initial individual choice and adopted the option selected by the partner. The dependent variable was therefore coded as 1 when a change of mind occurred and 0 otherwise. Because changes of mind only occurred in disagreement trials, we included only disagreement trials in this analysis. All candidate models included arbitrator’s own confidence as a predictor, operationalized as the confidence explicitly reported by the participant for their initial individual choice (discrete values ranging from 1 to 6). Partner’s encoded confidence was derived from the single-choice encoding model and corresponded to the posterior probability (values between 0 and 1) that the partner’s action encoded high confidence. To test whether partner’s encoded confidence influenced the probability of changes of mind, we compared a series of models of increasing complexity. We began with a basic model containing only arbitrator’s own confidence and no information about partner’s encoded confidence. We then added partner’s encoded confidence as an additional fixed effect. Finally, we fit a piece-wise model in which the effect of partner’s encoded confidence was allowed to vary across three levels of arbitrator’s own confidence (low, intermediate, and high). See Supplementary Tables S6a-d.

For each model, we included a participant-specific random intercept to account for stable interindividual differences in change propensity. No random slopes were included, as they did not improve model fit. Model performance was evaluated using information criteria (Bayesian Information Criterion) that penalize for complexity, enabling us to determine whether the added predictors and the piece-wise specification were justified after accounting for model complexity.

### Computational model

To examine how collective accuracy depends on the arbitrator’s integration of the partner’s confidence, we developed a computational model that systematically varied the weight given to the partner’s encoded confidence. We modelled the log-probability of changing one’s mind as a linear function of the arbitrator’s own confidence and the encoded partner’s confidence, replicating the fixed-effect structure of the GLMM fitted on empirical data. Probabilities predicted under different parameter configurations were then used to simulate single-trial choices and estimate collective accuracy under different behavioral scenarios. The simulation analysis was restricted to disagreement trials where the arbitrator’s own confidence level fell within the intermediate range, as these were the trials in which we empirically observed an effect of encoded confidence on the probability of change of mind. Specifically, the probability of changing one’s mind was simulated as:

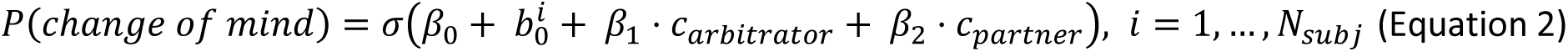

In the above equation, *σ* denotes the sigmoid function; *β*_0_ is the overall intercept of the model; *b_0_^i^* is a participant-specific random intercept capturing stable interindividual differences in baseline change propensity; *β*_1_ is the fixed-effect coefficient representing the influence of the arbitrator’s own confidence on change of mind; *β*_2_ is the fixed-effect coefficient representing the influence of the partner’s encoded confidence on change of mind; *c_arbitrator_* is the arbitrator’s own confidence at the single-trial level, and *c_partner_* is the partner’s encoded confidence at the single-trial level, both robustly z-scored.

To ground the simulation in empirically observed behavior, coefficients *β*_1_ and *b^i^* were assigned fixed values that corresponded to the coefficients of the GLMM fitted to the empirical data (see previous section and Table S6d). These fixed parameters preserved realistic individual differences in baseline change propensity (*b_0_^i^*) and the natural relationship between the arbitrator’s own confidence and their decisions to change their mind (*β*_1_).

In contrast, coefficients *β*_0_ and *β*_2_were varied systematically and beyond the range observed in real data to understand their effect on the probability of change of mind and model a broader range of possible behavioral scenarios (see Fig. 3C). Specifically:

− *β*_0_ controls the overall probability of changing one’s mind. We varied *β*_0_ from *β_EXP_* – 2.0 to *β*_0,*emp*_ + 2.0 in steps of 1.0, where *β*_0,*emp*_ is the intercept fitted to the empirical data. This manipulation allowed us to simulate scenarios where arbitrators were generally more or less willing to revise their initial decisions.

− *β*_2_ determines how strongly the partner’s confidence influences the arbitrator’s decision to change their mind. We varied *β*_2_ from –2.0 to 2.0 in steps of 0.05, a range that encompassed the empirically observed value of 0.593.

For each configuration of these parameters, we generated single-trial changes of mind (yes/no) as the most likely outcome given the value of the model’s probability of change of mind in that trial (that is we assigned it to a yes if the probability of change of mind was > 0.5 and to a no otherwise). We then computed collective accuracy based on the collective decisions derived from the simulated changes of mind in each scenario (see ‘deterministic model’, Tables S7 a-b). We also tested the alternative method of generating binomial choices with the probability expressed by the model in that trial, obtaining comparable results (see ‘stochastic model’, Tables S7 c-d)

The impact of varying *β*_0_ and *β*_2_ on the resulting collective accuracy was statistically assessed using GLMMs with a binomial family and a ‘logit’ link function, estimated collective accuracy (1=correct, 0=incorrect) as a dependent variable, *β*_0_ and *β*_2_ and their interaction as fixed effects, and participant as random intercept.

## Standardization of numerical variables

Numerical values for kinematic variables and for confidence ratings were standardized (z-scored) before analysis. Specifically, for kinematic variables, standard z-scores were computed, per participant, by first subtracting each value’s mean and then dividing by its standard deviation. Z-scored values were then used for training the kinematic encoding models. The z-scoring improves numerical stability during model training and helps prevent certain features from disproportionately influencing the classification due to different units or magnitudes.

For confidence ratings, robust z-scores were computed, per participant, by first subtracting each value’s median and then dividing by its median absolute deviation (MAD). Robust z-scoring was applied due to its robustness to non-normal distributions and outliers, as confidence ratings often exhibit skewed distributions or extreme values due to individual response biases (e.g., see Fig. S2). Z-scored values were then used in all abovementioned GLMMs that include reported confidence as factor. Values were z-scored per participant to account for the interindividual difference in how participants used the confidence scale. This way, the models were sensitive to relative differences in each participant’s confidence ratings rather than in absolute scale usage. Importantly, we verified that the classification of confidence ratings into three categories (described above; see Fig. S2) did not change regardless of whether it was applied to the raw values or to the z-scored values, indicating that the transformation preserved the original value distribution.

Robust z-scores were also computed for the posterior probabilities outputted by the encoding model. Here, we applied z-scoring across, rather than within, participants. This was done because encoding models can produce posterior probabilities with varying ranges and variances across participants due to differences in model fit quality or behavioral idiosyncrasies. Using unstandardized outputs in group-level regressions risks conflating these between-subject differences with true effects of interest. By z-scoring across participants, we standardized these distributions and reduced between-subject variance, resulting in more reliable and interpretable estimates of population-level effects. Z-scored values were then used in in all abovementioned GLMMs that include ‘encoded’ confidence as a factor.

In exploratory analyses, we confirmed that standard (mean-based) and robust (median-based) z-scoring yielded qualitatively similar results, suggesting that our findings are not sensitive to the specific choice of standardization method.

## Data, Materials, and Software Availability

The dataset is publicly available at the Open Science Framework (OSF) platform and can be accessed at the following link: https://doi.org/10.17605/OSF.IO/KE564. The code supporting the main results of this study is based on publicly available tools and has also been deposited on OSF.

## Acknowledgements

This work was supported by the European Commission under Horizon Europe, grant number 101092889, project SHARESPACE.

## Author contributions

LS, CB, AC, and BB conceived the study. LS, CB, AC and MM designed the experiment with input from BB. LS, CB, OS, NM, and SP designed the analyses. SP and NM conceived the computational model. LS and MM collected the data. LS, OS, and NM analyzed the data. LS and CB wrote the first draft of the manuscript with contributions from BB, SP, and NM. All authors revised and approved the final manuscript. CB supervised the work and acquired funding.

