## Supplemental Figures for "Confidence Read from Others’ Movements Guides Collective Decisions"

**Supplementary Figures**

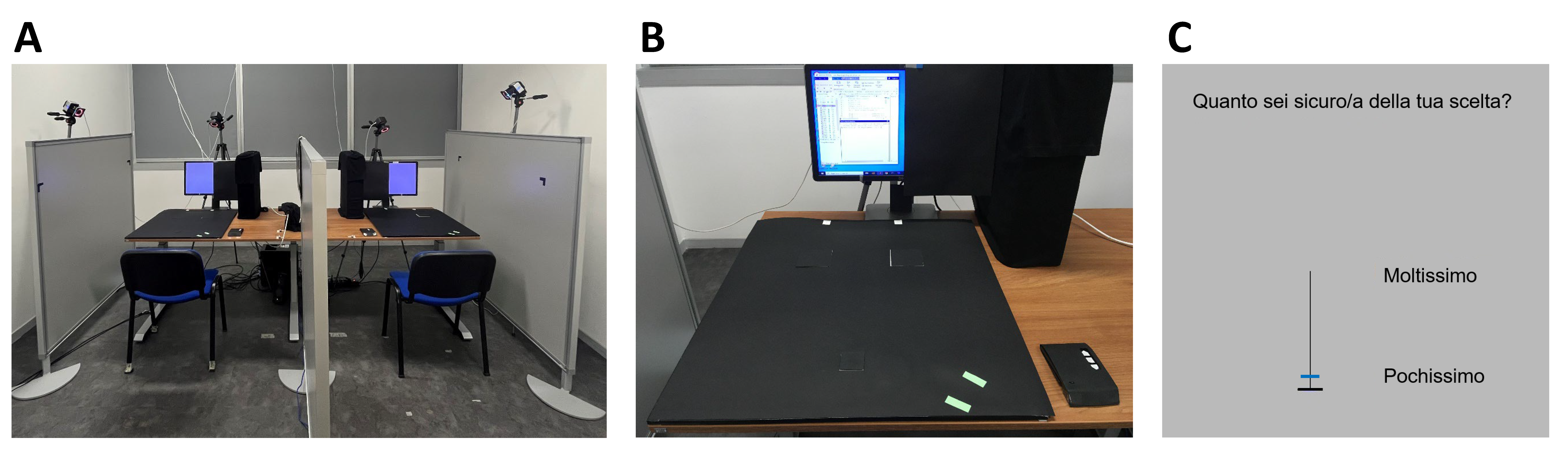

***Fig. S1 (related to Fig. 1A).*** ***(A) Setup.*** *Two participants (dyad members) were seated in front of individual computer screens, with a vertical divider positioned half-way in between. This divider prevented visual access between participants; they could see only the partner’s arm when it reached forward. Near-infrared cameras were positioned around the setup to record participants’ movement kinematics.* ***(B) Response panel.*** *The start position (small square) was located centrally in front of the participant and the two response positions (larger squares) were placed 36 cm away from the start position, offset by 12 cm to the right and left of the panel’s midline, respectively. After participants had responded, they privately rated how confident they felt in their choice using a keypad (shown on the right on the table).* ***(C) Confidence scale.*** *Participants rated how confident they felt in their choices by moving the slider up or down the scale (6 discrete levels, from 1 (very little; “pochissimo”) to 6 (very much; “moltissimo”) with the upper or lower arrow key on the keypad. To confirm their rating, participants pressed the middle key.*

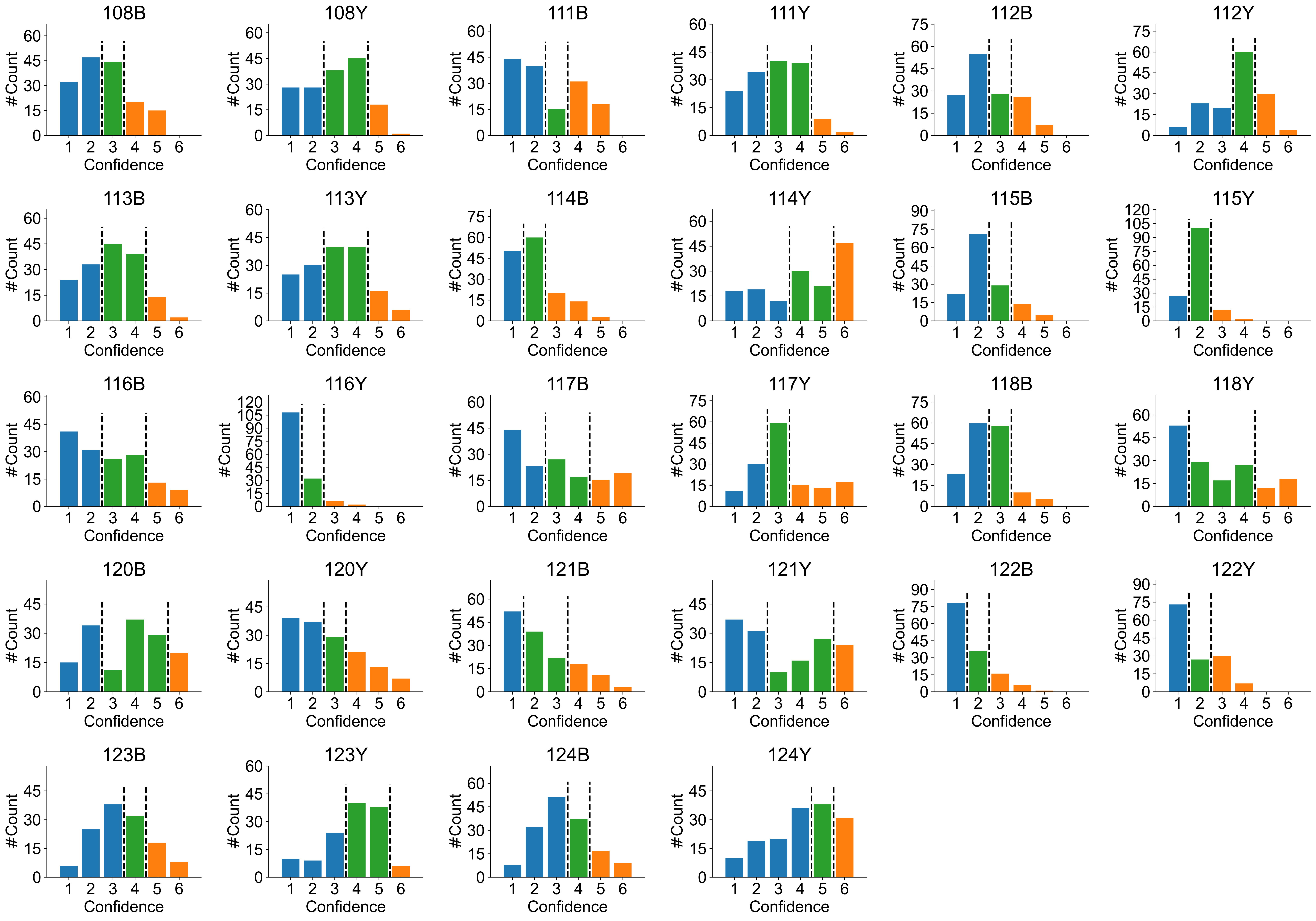

***Fig. S2. Categorizing individual confidence.*** *We categorized each participant’s self-reported confidence ratings (values ranging from 1 to 6) into three categories: Low (blue), Medium (green), and High (orange). For each participant, the confidence ratings from all individual choices (160 trials) were included. To categorize these ratings, we first identified, for each participant, two breakpoints corresponding to the 33^rd^ and 66^th^ percentiles of that participant’s confidence distribution. Values below or equal to the first breakpoint (≤ 33%) were labeled Low, values above the first and below or equal to the second breakpoint (> 33% and ≤ 66%) were labeled Medium, and values above the second breakpoint (> 66%) were labeled High. In rare instances (i.e., for 3 out of 28 participants), the two breakpoints coincided, splitting the distribution in only two instead of three categories. In these cases, we proceeded as follows. If the coinciding breakpoints were equal to the lowest confidence value of 1, we increased one breakpoint by 1. This resulted in the following categories: values of 1 corresponded to Low, values of 2 corresponded to Medium, and values from 3 to 6 corresponded to High (see participant 116Y). If the coinciding breakpoints were greater than 1, we decreased one breakpoint by 1. For instance, if the initial two breakpoints had the value 3, then one was transformed into 2, resulting in the following categories: values of 1-2 corresponded to Low, values of 3 corresponded to Medium, and values from 4 to 6 corresponded to High (see 117Y; see also 115Y for which the initial breakpoints equaled 2). Overall, this categorization procedure resulted in the following distribution across participants: Low: 398 trials (48.7%); Medium: 316 (38.6%); High = 104 (12.7%).*
