## Supplemental Tables for "Confidence Read from Others’ Movements Guides Collective Decisions"

**Supplementary Tables**

**Table S1.** (related to Fig. 1D)

Reported confidence predicts individual accuracy

| Probability of taking the correct decision | | | | |
| --- | --- | --- | --- | --- |
| Accuracy ~ Agreement * Confidence + (1 + Confidence \| Subject) | | | | |
| *Fixed effects* | *Estimate* | *SE* | *z* | *p* |
| (Intercept) | **0.317** | **0.109** | **2.902** | **0.004**** |
| Agreement | **1.098** | **0.120** | **9.180** | **<0.001***** |
| Confidence (H-M) | **0.431** | **0.207** | **2.082** | **0.037**** |
| Confidence (L-M) | **-0.621** | **0.122** | **-5.103** | **<0.001***** |
| Agreement : Confidence (H-M) | 0.376 | 0.244 | 1.543 | 0.123 |
| Agreement : Confidence (L-M) | 0.038 | 0.154 | 0.248 | 0.804 |

| EMMs of Accuracy by Agreement and Confidence level | | |
| --- | --- | --- |
| *Agreement* |  | |
| *Confidence* | *Estimate* | *SE* |
| Low | 0.697 | 0.015 |
| Medium | 0.805 | 0.017 |
| High | 0.902 | 0.019 |
| *Disagreement* |  |  |
| *Confidence* | *Estimate* | *SE* |
| Low | 0.425 | 0.018 |
| Medium | 0.579 | 0.027 |
| High | 0.679 | 0.049 |

*Note:* Reported confidence values have been standardized and categorized into three levels (L=Low, M=Medium, H=High) within participants. For the EMMs, results are reported separately for agreement vs. disagreement trials. Intervals are back-transformed from the logit scale.

**Table S2a.** (related to Fig. 1C, left panel)

Perceptual sensitivity for individual vs. collective decisions (all participants)

| Psychometric curve (probability of reporting 2^nd^ interval) | | | | |
| --- | --- | --- | --- | --- |
| P(2^nd^ interval) ~ Decision type * Contrast level + (1 + Contrast level \| Subject) | | | | |
| *Fixed effects* | *Estimate* | *SE* | *z* | *p* |
| (Intercept) | **-0.208** | **0.058** | **-3.612** | **<0.001***** |
| Decision type | -0.018 | 0.041 | -0.431 | 0.666 |
| Contrast level | **8.527** | **0.855** | **9.972** | **<0.001***** |
| Decision type : Contrast level | **1.706** | **0.566** | **3.011** | **0.003**** |

| Simple psychometric slopes by Decision type | | | | |  | |
| --- | --- | --- | --- | --- | --- | --- |
| *Decision type* | *Estimate* | *SE* | *df* | *z* | | *p* |
| Individual | **8.527** | **0.855** | **Inf** | **9.972** | | **<0.001***** |
| Collective | **10.233** | **0.873** | **Inf** | **11.720** | | **<0.001***** |

| Psychometric slope difference: individual vs. collective decisions | | | |  | | |
| --- | --- | --- | --- | --- | --- | --- |
| *Contrast* | *Estimate* | *SE* | *df* | | *z* | *p* |
| Individual - Collective | **-1.706** | **0.566** | **Inf** | | **-3.011** | **0.003**** |

**Table S2b.** (related to Fig. 1C, middle panel)

Perceptual sensitivity for individual vs. collective decisions (“min” participants)

| Psychometric curve (probability of reporting 2^nd^ interval) | | | | |
| --- | --- | --- | --- | --- |
| P(2^nd^ interval) ~ Decision type * Contrast level + (1 + Contrast level \| Subject) | | | | |
| *Fixed effects* | *Estimate* | *SE* | *z* | *p* |
| (Intercept) | **-0.211** | **0.081** | **-2.609** | **0.009**** |
| Decision type | -0.060 | 0.058 | -1.039 | 0.299 |
| Contrast level | **7.160** | **1.169** | **6.127** | **<0.001***** |
| Decision type : Contrast level | **3.417** | **0.794** | **4.301** | **<0.001***** |

| Simple psychometric slopes by Decision type | | | | | |
| --- | --- | --- | --- | --- | --- |
| *Decision type* | *Estimate* | *SE* | *df* | *z* | *p* |
| Individual | **7.160** | **1.169** | **Inf** | **6.127** | **<0.001***** |
| Collective | **10.576** | **1.216** | **Inf** | **8.699** | **<0.001***** |

| Psychometric slope difference: Individual vs. collective decisions | | | | | |
| --- | --- | --- | --- | --- | --- |
| *Contrast* | *Estimate* | *SE* | *df* | *z* | *p* |
| Individual - Collective | **-3.417** | **0.794** | **Inf** | **-4.301** | **<0.001***** |

**Table S2c.** (related to Fig. 1C, right panel)

Perceptual sensitivity for individual vs. collective decisions (“max” participants)

| Psychometric curve (probability of reporting 2^nd^ interval) | | | | |
| --- | --- | --- | --- | --- |
| P(2^nd^ interval) ~ Decision type * Contrast level + (1 + Contrast level \| Subject) | | | | |
| *Fixed effects* | *Estimate* | *SE* | *z* | *p* |
| (Intercept) | **-0.208** | **0.082** | **-2.530** | **0.011*** |
| Decision type | -0.024 | 0.058 | -0.431 | 0.680 |
| Contrast level | **10.051** | **1.246** | **8.066** | **<0.001***** |
| Decision type : Contrast level | -0.147 | 0.819 | -0.179 | 0.858 |

**Table S3a.** (related to Fig. 1E, left panel)

Accuracy for individual vs. collective decisions (all participants)

| Probability of taking the correct decision | | | | |
| --- | --- | --- | --- | --- |
| Accuracy ~ Decision type * Contrast level + (1 + Contrast level \| Subject) | | | | |
| *Fixed effects* | *Estimate* | *SE* | *z* | *p* |
| (Intercept) | -0.102 | 0.081 | -1.263 | 0.207 |
| Decision type | -0.020 | 0.103 | -0.198 | 0.843 |
| Contrast level | **14.536** | **1.611** | **9.025** | **<0.001***** |
| Decision type : Contrast level | **3.000** | **1.509** | **1.988** | **0.047*** |

| EMMs of Accuracy by Contrast level and Decision type | | | |
| --- | --- | --- | --- |
| *Contrast level* | *Decision type* | *Estimate* | *SE* |
| *0.015* | Individual | 0.529 | 0.018 |
| *0.015* | Collective | 0.535 | 0.018 |
| *0.035* | Individual | 0.600 | 0.017 |
| *0.035* | Collective | 0.620 | 0.017 |
| *0.070* | Individual | 0.714 | 0.021 |
| *0.070* | Collective | 0.751 | 0.019 |
| *0.150* | Individual | 0.889 | 0.021 |
| *0.150* | Collective | 0.925 | 0.015 |

| Pairwise comparisons of Decision type (Individual vs. Collective) at each Contrast level | | | | | |
| --- | --- | --- | --- | --- | --- |
| *Contrast level* | *Estimate* | *SE* | *df* | *z* | *p* |
| 0.015 | -0.006 | 0.022 | Inf | -0.286 | 0.775 |
| 0.035 | -0.020 | 0.017 | Inf | -1.200 | 0.230 |
| 0.070 | **-0.037** | **0.014** | **Inf** | **-2.666** | **0.008**** |
| 0.150 | **-0.036** | **0.015** | **Inf** | **-2.473** | **0.013*** |

*Note:* For EMMs, intervals are back-transformed from the logit scale. Pairwise comparisons were performed using Tukey-adjusted tests.

**Table S3b.** (related to Fig. 1E, middle panel)

Accuracy for individual vs. collective decisions (“min” participants)

| Probability of taking the correct decision | | | | |
| --- | --- | --- | --- | --- |
| Accuracy ~ Decision type * Contrast level + (1 \| Subject) | | | | |
| *Fixed effects* | *Estimate* | *SE* | *z* | *p* |
| (Intercept) | 0.022 | 0.141 | 0.157 | 0.876 |
| Decision type | 0.029 | 0.146 | 0.196 | 0.844 |
| Contrast level | **10.029** | **1.343** | **7.465** | **<0.001***** |
| Decision type : Contrast level | **4.809** | **2.092** | **2.299** | **0.022*** |

| EMMs of Accuracy by Contrast level and Decision type | | | |
| --- | --- | --- | --- |
| *Contrast level* | *Decision type* | *Estimate* | *SE* |
| *0.015* | Individual | 0.543 | 0.033 |
| *0.015* | Collective | 0.568 | 0.033 |
| *0.035* | Individual | 0.592 | 0.029 |
| *0.035* | Collective | 0.639 | 0.028 |
| *0.070* | Individual | 0.674 | 0.026 |
| *0.070* | Collective | 0.748 | 0.023 |
| *0.150* | Individual | 0.821 | 0.025 |
| *0.150* | Collective | 0.907 | 0.017 |

| Pairwise comparisons of Decision type (Individual vs. Collective) at each Contrast level | | | | | |
| --- | --- | --- | --- | --- | --- |
| *Contrast level* | *Estimate* | *SE* | *df* | *z* | *p* |
| 0.015 | -0.025 | 0.031 | Inf | -0.816 | 0.414 |
| 0.035 | -0.047 | 0.024 | Inf | -1.954 | 0.0507 |
| 0.070 | **-0.075** | **0.020** | **Inf** | **-3.707** | **<0.001***** |
| 0.150 | **-0.085** | **0.026** | **Inf** | **-3.328** | **<0.001***** |

*Note:* For EMMs, intervals are back-transformed from the logit scale. Pairwise comparisons were performed using Tukey-adjusted tests.

**Table S3c.** (related to Fig. 1E, right panel)

Accuracy for individual vs. collective decisions (“max” participants)

| Probability of taking the correct decision | | | | |
| --- | --- | --- | --- | --- |
| Accuracy ~ Decision type * Contrast level + (1 \| Subject) | | | | |
| *Fixed effects* | *Estimate* | *SE* | *z* | *p* |
| (Intercept) | -0.124 | 0.134 | -0.928 | 0.354 |
| Decision type | -0.039 | 0.148 | -0.261 | 0.794 |
| Contrast level | **15.959** | **1.596** | **9.998** | **<0.001***** |
| Decision type : Contrast level | 0.243 | 2.252 | 0.108 | 0.914 |

**Table S4a.**

Arbitrator’s own confidence predicts changes of mind (all trials)

| Probability of Change of Mind | | | | |
| --- | --- | --- | --- | --- |
| Change of Mind ~ Confidence + (1 \| Subject) | | | | |
| *Fixed effects* | *Estimate* | *SE* | *z* | *p* |
| (Intercept) | **-1.899** | **0.153** | **-12.386** | **<0.001***** |
| Confidence (High-Medium) | **-1.178** | **0.254** | **-4.633** | **<0.001***** |
| Confidence (Low-Medium) | **1.290** | **0.133** | **9.684** | **<0.001***** |

| EMMs of Change of Mind probability by Confidence level | | |
| --- | --- | --- |
| *Confidence level* | *Estimate* | *SE* |
| Low | 0.352 | 0.030 |
| Medium | 0.130 | 0.017 |
| High | 0.044 | 0.011 |

| Pairwise comparisons of Change of Mind probability between Confidence levels | | | | | |
| --- | --- | --- | --- | --- | --- |
| *Confidence level* | *Estimate* | *SE* | *df* | *z* | *p* |
| Low-Medium | **0.222** | **0.025** | **Inf** | **9.032** | **<0.001***** |
| Low-High | **0.308** | **0.028** | **Inf** | **10.970** | **<0.001***** |
| Medium-High | **0.086** | **0.017** | **Inf** | **4.968** | **<0.001***** |

*Note:* For EMMs, intervals are back-transformed from the logit scale. Pairwise comparisons were performed using Tukey-adjusted tests.

**Table S4b.** (related to Fig. 1F)

Arbitrator’s own confidence predicts changes of mind (disagreement trials only)

| Probability of change of mind | | | | |
| --- | --- | --- | --- | --- |
| Change of Mind ~ Confidence + (1 \| Subject) | | | | |
| *Fixed effects* | *Estimate* | *SE* | *z* | *p* |
| (Intercept) | **-0.886** | **0.243** | **-3.654** | **<0.001***** |
| Confidence (High-Medium) | **-1.092** | **0.314** | **-3.474** | **<0.001***** |
| Confidence (Low-Medium) | **2.440** | **0.219** | **11.163** | **<0.001***** |

| EMMs of Change of Mind by Confidence level | | |
| --- | --- | --- |
| *Confidence level* | *Estimate* | *SE* |
| Low | 0.826 | 0.035 |
| Medium | 0.292 | 0.050 |
| High | 0.122 | 0.038 |

| Pairwise comparisons of Change of Mind probability between Confidence levels | | | | | |
| --- | --- | --- | --- | --- | --- |
| *Confidence level* | *Estimate* | *SE* | *df* | *z* | *p* |
| Low-Medium | **0.534** | **0.041** | **Inf** | **13.111** | **<0.001***** |
| Low-High | **0.704** | **0.040** | **Inf** | **17.639** | **<0.001***** |
| Medium-High | **0.170** | **0.046** | **Inf** | **3.739** | **<0.001***** |

*Note:* For EMMs, intervals are back-transformed from the logit scale. Pairwise comparisons were performed using Tukey-adjusted tests.

**Table S5.** (related to Fig. 2D)

Cross-validated accuracy of encoding models

| Results of Stratified K-Fold Cross-Validation(100 x 5-fold, 1000 repetitions) per Subject | | | | | |
| --- | --- | --- | --- | --- | --- |
| *Subject* | *Trials* | *Low confidence choices* | *High confidence choices* | *Balanced Accuracy* | *p-value* |
| 01P1 | 158 | 79 | 79 | 0,701 | **<0.001** |
| 01P2 | 158 | 94 | 64 | 0,544 | 0.189 |
| 02P1 | 148 | 84 | 64 | 0,850 | **<0.001** |
| 02P2 | 148 | 58 | 90 | 0,821 | **<0.001** |
| 03P1 | 143 | 82 | 61 | 0,599 | **0.023** |
| 03P2 | 143 | 49 | 94 | 0,571 | 0.063 |
| 04P1 | 157 | 57 | 100 | 0,600 | **0.022** |
| 04P2 | 157 | 95 | 62 | 0,616 | **0.006** |
| 05P1 | 147 | 50 | 97 | 0,671 | **<0.001** |
| 05P2 | 147 | 79 | 68 | 0,650 | **0.002** |
| 06P1 | 141 | 93 | 48 | 0,617 | **0.014** |
| 06P2 | 141 | 27 | 114 | 0,465 | 0.735 |
| 07P1 | 148 | 72 | 76 | 0,776 | **<0.001** |
| 07P2 | 148 | 108 | 40 | 0,725 | **<0.001** |
| 08P1 | 145 | 67 | 78 | 0,565 | 0.068 |
| 08P2 | 145 | 100 | 45 | 0,766 | **<0.001** |
| 09P1 | 156 | 83 | 73 | 0,670 | **<0.001** |
| 09P2 | 156 | 82 | 74 | 0,734 | **<0.001** |
| 10P1 | 146 | 60 | 86 | 0,794 | **<0.001** |
| 10P2 | 146 | 76 | 70 | 0,756 | **<0.001** |
| 11P1 | 145 | 91 | 54 | 0,835 | **<0.001** |
| 11P2 | 145 | 68 | 77 | 0,790 | **<0.001** |
| 12P1 | 137 | 78 | 59 | 0,727 | **<0.001** |
| 12P2 | 137 | 73 | 64 | 0,623 | **0.005** |
| 13P1 | 127 | 69 | 58 | 0,519 | 0.330 |
| 13P2 | 127 | 83 | 44 | 0,652 | **0.002** |
| 14P1 | 154 | 91 | 63 | 0,538 | 0.176 |
| 14P2 | 154 | 85 | 69 | 0,741 | **<0.001** |

*Note:* The first column of the table contains participant IDs, consisting of pair number (01-14) and participant number (P1 and P2). The second column indicates the number of available trials for each pair, while columns 3-4 indicate the number of choices each participant made with low and high confidence, respectively (a median split was used to determine the two categories low vs. high per participant). Column 5 shows the performance (in terms of Balanced Accuracy) of each participant’s model, assessed using repeated stratified 5-fold cross-validation with 100 random splits. The last column indicates whether the model performed significantly above chance when estimating the reported confidence from the kinematics.

**Table S6a.**

Model 0: No partner confidence

| Probability of change of mind | | | | |
| --- | --- | --- | --- | --- |
| P(change) ~ Arbitrator Confidence + (1 \| Subject) | | | | |
| *Fixed effects* | *Estimate* | *SE* | *z* | *p* |
| (Intercept) | 0.274 | 0.208 | 1.320 | 0.187 |
| Arbitrator Confidence | **-1.804** | **0.148** | **-12.203** | **<0.001***** |

**Table S6b.**

Model 1: Partner’s reported confidence

| Probability of change of mind | | | | |
| --- | --- | --- | --- | --- |
| P(change) ~ Arbitrator Confidence + Partner Rep Confidence + (1 \| Subject) | | | | |
| *Fixed effects* | *Estimate* | *SE* | *z* | *p* |
| (Intercept) | 0.282 | 0.205 | 1.373 | 0.170 |
| Arbitrator Confidence | **-1.793** | **0.148** | **-12.122** | **<0.001***** |
| Partner Rep Confidence | 0.130 | 0.107 | 1.220 | 0.223 |

**Table S6c.**

Model 2: Partner’s encoded confidence

| Probability of change of mind | | | | |
| --- | --- | --- | --- | --- |
| P(change) ~ Arbitrator Confidence + Partner Enc Confidence + (1 \| Subject) | | | | |
| *Fixed effects* | *Estimate* | *SE* | *z* | *p* |
| (Intercept) | 0.323 | 0.205 | 1.574 | 0.115 |
| Arbitrator Confidence | **-1.792** | **0.148** | **-12.131** | **<0.001***** |
| Partner Enc Confidence | **0.249** | **0.105** | **2.372** | **0.018*** |

**Table S6d.** (related to Fig. 3A)

Model 3: Piece-wise model: Partner’s encoded confidence varied across arbitrator’s confidence

| Probability of change of mind | | | | |
| --- | --- | --- | --- | --- |
| P(change) ~ Arbitrator Confidence + Partner Enc Confidence : Arbitrator Confidence [L,M,H] + (1 \| Subject) | | | | |
| *Fixed effects* | *Estimate* | *SE* | *z* | *p* |
| (Intercept) | 0.318 | 0.205 | 1.553 | 0.120 |
| Arbitrator Confidence | **-1.762** | **0.152** | **-11.600** | **<0.001***** |
| Partner Enc Conf : Arbitrator Conf M | **0.593** | **0.172** | **3.450** | **<0.001***** |
| Partner Enc Conf : Arbitrator Conf L | 0.045 | 0.141 | 0.322 | 0.748 |
| Partner Enc Conf : Arbitrator Conf H | -0.071 | 0.392 | -0.182 | 0.856 |

**Table S6e.**

Goodness-of-fit and model comparisons

| *Model* | *Fixed effects* | *df* | *BIC* | *Deviance* | *p (LRT)* |
| --- | --- | --- | --- | --- | --- |
| M0 | Arbitrator Confidence | 3 | 876.5 | 856.4 |  |
| M1 | + Partner Reported Confidence | 4 | 881.7 | 854.9 | vs. M0: 0.223 |
| M2 | + Partner Encoded Confidence | 4 | 877.6 | 850.7 | vs. M0: **0.017** |
| M3 | + Partner Enc Conf : Arbitrator Conf [L,M,H] | 6 | 883.8 | 843.6 | vs. M2: **0.028** |

*Note:* Likelihood-ratio tests (LRTs) were used for nested model comparisons. The comparison between M1 and M2 relied on Bayesian Information Criterion (BIC) because the two models include the same number of parameters; the lower BIC indicates a preferable fit for M2.

**Table S7a.** (related to Fig. 3B)

Deterministic model: Effects of baseline change tendency ($\beta_{0}$) and partner encoded confidence ($\beta_{2}$) on simulated collective accuracy

| *Coefficient* | *Estimate* | *SE* | *95% CI* | *z* | *p* |
| --- | --- | --- | --- | --- | --- |
| (Intercept) | 0.1172 | 0.0634 | –0.0115, 0.2459 | 1.849 | 0.0645 |
| $\boldsymbol{\beta}_{\mathbf{2}}$ | **0.0817** | **0.0042** | **0.0736, 0.0899** | **19.629** | **<0.0001** |
| $\boldsymbol{\beta}_{\mathbf{0}}$ | **–0.1401** | **0.0039** | **–0.1477, –0.1325** | **–36.008** | **<0.0001** |
| $\boldsymbol{\beta}_{\mathbf{0}}$ **:** $\boldsymbol{\beta}_{\mathbf{2}}$ | **0.0195** | **0.0030** | **0.0137, 0.0253** | **6.595** | **<0.0001** |

*Note:* The GLMM includes simulated collective accuracy (1=correct, 0=incorrect) as a dependent variable, and β_0_, β_2_, and their interaction as fixed effects, and a participant-specific random intercept.

**Table S7b.** (related to Fig. 3D)

Deterministic model: Effect of partner encoded confidence ($\beta_{2}$) on simulated collective accuracy across baseline levels ($\beta_{0}$)

| $\beta_{0}$ | *Slope* | *SE* | *95% CI* | *z* | *p* |
| --- | --- | --- | --- | --- | --- |
| $\boldsymbol{\beta}_{\mathbf{0,emp}}$ **­– 2.0** | **0.0428** | **0.0072** | **0.0241, 0.0614** | **5.911** | **<0.0001** |
| $\boldsymbol{\beta}_{\mathbf{0,emp}}$ **­– 1.0** | **0.0622** | **0.0051** | **0.0491, 0.0754** | **12.177** | **<0.0001** |
| $\boldsymbol{\beta}_{\mathbf{0,emp}}$ | **0.0887** | **0.0042** | **0.0710, 0.0924** | **19.629** | **<0.0001** |
| $\boldsymbol{\beta}_{\mathbf{0}\mathbf{,emp}}$ **­+ 1.0** | **0.1012** | **0.0051** | **0.0881, 0.1143** | **19.856** | **<0.0001** |
| $\boldsymbol{\beta}_{\mathbf{0,emp}}$ **­+ 2.0** | **0.1207** | **0.0072** | **0.1021, 0.1392** | **16.725** | **<0.0001** |

**Table S7c.**

Stochastic model: Effects of baseline change tendency ($\beta_{0}$) and partner encoded confidence ($\beta_{2}$) on simulated collective accuracy

| *Coefficient* | *Estimate* | *SE* | *95% CI* | *z* | *p* |
| --- | --- | --- | --- | --- | --- |
| **(Intercept)** | **0.0971** | **0.0495** | **–0.0032, 0.1974** | **1.962** | **0.0498** |
| $\boldsymbol{\beta}_{\mathbf{2}}$ | **0.0607** | **0.0041** | **0.0526, 0.0688** | **14.695** | **<0.0001** |
| $\boldsymbol{\beta}_{\mathbf{0}}$ | **–0.0997** | **0.0039** | **–0.1073, –0.0922** | **–25.873** | **<0.0001** |
| $\boldsymbol{\beta}_{\mathbf{0}}$ **:** $\boldsymbol{\beta}_{\mathbf{2}}$ | **0.0128** | **0.0029** | **0.0071, 0.0185** | **4.359** | **<0.0001** |

**Table S7d.**

Stochastic model: Effect of partner encoded confidence ($\beta_{2}$) on simulated collective accuracy across baseline levels ($\beta_{0}$)

| $\beta_{0}$ | *Slope* | *SE* | *95% CI* | *z* | *p* |
| --- | --- | --- | --- | --- | --- |
| $\boldsymbol{\beta}_{\mathbf{0,emp}}$ **­– 2.0** | **0.0352** | **0.0072** | **0.0167, 0.0537** | **4.908** | **<0.0001** |
| $\boldsymbol{\beta}_{\mathbf{0,emp}}$ **­– 1.0** | **0.0479** | **0.0051** | **0.0349, 0.0610** | **9.460** | **<0.0001** |
| $\boldsymbol{\beta}_{\mathbf{0,emp}}$ | **0.0607** | **0.0041** | **0.0501, 0.0713** | **14.695** | **<0.0001** |
| $\boldsymbol{\beta}_{\mathbf{0,emp}}$ **­+ 1.0** | **0.0735** | **0.0051** | **0.0604, 0.0865** | **14.531** | **<0.0001** |
| $\boldsymbol{\beta}_{\mathbf{0,emp}}$ **­+ 2.0** | **0.0862** | **0.0072** | **0.0678, 0.1046** | **12.052** | **<0.0001** |
